# A deep learning-based automated closed-loop optogenetic system for neuromodulation during seizures

**DOI:** 10.1101/2022.04.11.487951

**Authors:** Tamal Batabyal, Anastasia Brodovskaya, John Williamson, Jaideep Kapur

## Abstract

Closed-loop electrical brain stimulation is becoming a popular technique proposed for use as a treatment alternative to surgical resection of brain tissue for drug-resistant seizures in epilepsy patients. Closed-loop optogenetic stimulation is an experimental alternative to electrical stimulation since it can stimulate or inhibit neurons. The closed-loop part contains an online seizure detection algorithm, which, based on the design, can detect the onset of a seizure or evaluate a running window for seizure membership. Conventional configurations of closed-loop optogenetics have several limitations, ranging from the adaptability of hardware-based implementation to inadequate, customized feature selection of seizures, among others. Here we provide a detailed description of our closed-loop components. We used a sequential, fully convolutional neural network regressor for complex feature selection of seizures against the controls from local field potential recordings. Our modular design kept the local field potential-recording headset and optical probe separate. The seizure detection and execution of light delivery are fast and can be precisely timed. This automated system is robust to noise, modular in design, flexible in use, and simple execution. When applied in vivo, the proposed work shows efficacy over the state of the arts in terms of improved seizure detection and reduction in false positives.

## 1. Introduction

Optogenetics is a recent technology that offers precise spatial, temporal, and cell-specific neuromodulation ^1–4^. It uses exogenous light-sensitive proteins called opsins, delivered using viral vectors to targeted brain regions. The specific neuronal population is accessed using different viral serotypes, enhancers, cell-type-specific promoters, and Cre recombinase techniques. Optogenetics can manipulate neural states and plasticity, control behaviors, identify micro and mesoscale circuits, and leverage precise access. In clinical settings, optogenetics is among potential therapeutic alternatives for neurological diseases, such as epilepsy and Parkinson’s disease, and can be used in cochlear, retinal, and motor prosthetics ^1,2,5–9^.

Optogenetics provides several advantages over state-of-the-art techniques. For example, conventional treatments in epilepsy, such as drugs, and surgical resection of epileptogenic foci, do not have spatiotemporal precision or cell-type specificity. These treatments cause adverse side effects. In a non-reversible treatment strategy, the surgical approaches need a well-defined, precisely located seizure focus. Faulty resection of healthy brain tissue can have debilitating consequences in patients. Furthermore, neocortical seizures can still remain refractory to such treatments ^10^. Optogenetics is a reversible tool, no brain tissue is permanently resected, and it offers spatiotemporally precise neuromodulation. Another well-known tool is electrical deep brain stimulation, a unipolar modulation ^11^, whereas optogenetics offers reversible excitation and inhibition of cellular activity based on the selected opsins.

In closed-loop optogenetic settings, simultaneous readout of ongoing neuronal activity is used to decide when and how to administer light. The term ‘closed loop’ originates from the control theory, where a transformation of the difference between the model output and the desired output is added to the input in a feedback path. This stereotypical model is widely used in electrical and computer engineering fields, such as oscillator design, adaptive filtering, and machine learning^12–15^. In neuroscience, closed-loop systems are real-time instantaneous feedback types ^2,16^ or input adaptation types ^17–19^. Problems such as seizure interruption fall into the first type. The second type can control neuronal firing rate. This work focuses on the closed-loop optogenetic system configurations used to detect and modulate seizures.

Many challenges in the design of a closed-loop system, including the complex non-linear pattern of electroencephalographic (EEG) or local field potential (LFP) data, variability in baseline over time, and non-linear noise, limit the identification of desired activity. Several artifacts from movement, muscle, and high-frequency sampling, are ubiquitous in EEG or LFP data. Linear filtering techniques cannot eliminate noise patterns that mimic EEG or LFP signals. Typical EEG or LFP signal (for example, K complex, sleep spindle, and seizure) correlated with a biological event (for example, sleep and epilepsy) also contains variation in the signal’s characteristics, such as the morphology amplitude and frequency. Therefore, a biological event/phenomenon classification using the EEG or LFP data needs a robustly-trained machine learning model.

The central part of closed-loop optogenetic settings for seizure interruption is the online seizure detection using LFP/EEG data. The detection and prediction of a seizure encompass a class of problems – seizure onset detection, seizure detection, and detection of seizure termination. These problems are binary classification tasks, where the data at an instant or a window of data (LFP/EEG) is tagged as 1 (seizure/seizure onset/ seizure termination) or 0 (otherwise). The strategy for light delivery in optogenetics depends on the users’ choice of classification task at hand. Generally speaking, there are two types of strategy (**Type 1**.) If a seizure onset is detected, light can be delivered *continuously* for a brief period (light ON) of time, followed by an inactivation period (light OFF) of delivery ^2^. No classification task is performed during the light delivery, which reduces computational overhead. However, this configuration is agnostic for neural activity during light delivery and seizure termination. Instead of seizure onset, an algorithm can evaluate a running data window (**Type 2**). An optogenetic driver can deliver light if the data in the window corresponds to the ongoing seizure. This setting does not require a manually-set inactivation routine. The algorithm assesses the neuronal activity on the fly and can stop the light delivery once the desired neuronal activity disappears. This approach seems suitable in instances where neuronal activity and cellular physiology are vulnerable to more prolonged light exposure. However, the inability of the configuration to concurrently access the data and trigger the light precludes continuous modulation of light delivery.

Apart from classification, there are implementation challenges. For a reliable, fully-automated, online optogenetics system, the concurrent execution of system modules should meet many requirements, such as the fast EEG or LFP acquisition, without interrupting the continuous recording and prompt evaluation of the acquired data light pulse probing, and others. Failure to comply with the requirements can have adverse consequences, such as alteration of cellular physiology and cell death.

In this work, we recorded in vivo neuronal LFP using recording software called Labchart. Using LFP, we proposed a configuration of an efficient, automated closed-loop system for optogenetic stimulation.

Our main contributions are:

a. We employed a deep learning-based regressor that extracts complex features of LFP recordings. The regressor identified and scored ongoing seizures and efficiently ignored non-seizure LFP. We did not customize features.
b. The system is fast and fully software controlled. Apart from the optogenetic toolbox and Arduino TTL pulse generator board, the system did not require external hardware like a digital signal processor.
c. Users can design the desired pattern of pulses in MATLAB.
d. In our design, independent recording LFP using the headset 16 and the optical probe, including the optogenetic system. Headset containing electrodes and optical probe are implanted separately. It offers flexibility in terms of customization of electrodes for preferred brain regions. We demonstrated in vivo the efficacy of our proposed configuration.

## 2. Literature Review

### 2.1 Seizure / seizure onset detection algorithms

Online feature selection of seizures is the central part of a closed-loop optogenetic system for seizure interruption. Osorio et al. used a wavelet FIR filter to perform automatic, real-time detection of seizures from EEG ^20,21^. In ^22^, the authors extracted features such as average half-wave duration, amplitude, dominant frequency, average power, spatial features from a multichannel EEG recording and used the nearest neighborhood classifier to predict the onset of a seizure. Based on the fractal dimension of a curve as defined by Katz ^23^, “line length” was proposed as a prominent feature of seizure onset ^24,25^ and it was used in ^2^. The algorithm to compute “line length” contains a subject-specific tunable parameter and the delay of electrographic seizure onset detection depends on such parameters, which is a drawback of this approach. High-end binary classifiers, such as a support vector machine, were used in studies where the features are customized using EEG data ^26–32^. Although these methods showed improved performance, a latency (2∼8 s) existed between the expert-marked and algorithm-detected seizure onset ^31^. This significant and variable latency will pose problems in clinical diagnosis and interruption of an ongoing seizure by light delivery. For example, a clinically meaningful seizure can be as short as10s ^33,34^. In this case, a latency of 4s in detecting the onset of a seizure will critically delay intervention.

Studies compared commercially available seizure detection software, such as Persyst 13, Encevis 1.7, and Besa 2.0. The detection rate varies from 67.6%-81% ^35^, substantially smaller than the reported accuracy (∼95-100%). In addition, the latency of detection is ℴ(10*s*). There are various sources of such anomaly – training data overfit in machine learning models used in research articles, computational demand (such as principal component analysis, complex transformations), a small corpus of training data, and data imbalance.

Deep learning-based approaches, such as convolutional neural networks (CNN), recurrent neural networks (RNN), and long short-term memory (LSTM) networks, are potential alternatives in seizure or seizure onset detection ^36,37^. Deep learning configurations are superior to the state-of-the-art machine learning models in many fields, ranging from object and activity detection ^38–40^ to language modeling ^41^ to biological problems ^42–44^. If properly trained with a large corpus of data, a deep learning model can extract salient features of the complex polymorphic pattern of a seizure. In addition, it provides a faster prediction with improved performance compared to state-of-the-art methods. It is difficult to compare all the deep learning configurations with our methods. These methods are trained on various datasets, such as raw EEG traces, multichannel EEG, pre-transformed (such as wavelet, fast Fourier and Stockwell transform) EEG traces, EEG periodograms, EEG power spectra, and in conjunction with other data modality like ECG. Furthermore, many such methods are for offline instead of online seizure detection. The majority of these studies considered EEG data instead of LFP data. These deep learning models can be easily deployed for seizure or seizure onset detection upon training with LFP data.

We used a deep, 1D fully convolutional feedforward neural regressor network to score a fixed-length running window of LFP data. The reason for the choice of CNN is two-fold. Firstly, in a previous study by ^45^, it is reported that CNN outperforms RNN and fully connected neural network (FCNN) in seizure detection using EEG data. This study was reproduced in ^46^ with an empirical observation that 1DCNN performs elegantly for seizure detection using EEG data. Secondly, there are a lot of seizure detection models that use LSTM in the literature ^37,47–49^. LSTM offers an elegant choice for seizure classification for time-series data that can exploit the hidden relationship between currently acquired data with the one at previous instants. However, the LFP data acquisition in our case is not continuous. We interleave data acquisition and the light triggering sequentially, creating a “silent” period (light delivery) between two consecutive instants of data acquisition. The barrier to performing them both in parallel is that two different programs can not simultaneously access LabChart data. Additionally, the accessed data at an instant for some time (let’s say 50 ms at 10kHz) is of variable (405 ± 90) length due to the stochasticity of peripheral communication between MATLAB and LabChart. This may alter the hidden relationship in data and cause problems while training an LSTM. So, we resorted to 1D CNN.

### 2.2 Closed-loop optogenetic configurations for in vivo seizure interruption

Several ways to design a closed-loop optogenetic system in the literature involve neurobiological experiments on several areas, such as learning, memory, and seizure. Authors in ^50^ devised a multichannel LFP recording system, where the command signal for optogenetic light delivery was based on EEG spike detection in prototypical settings. The spikes were sorted using a template matching algorithm, where the template was constructed by applying the EM algorithm to the baseline LFP. Although the configuration is fast in execution and can afford LFP acquisition at an excellent spatial precision, the spike sorting would fail in real-time to distinguish seizures because of the variations in baseline LFP and complex spike waveforms during seizures. Authors ^51,52^ proposed a closed-loop optogenetic system that can rapidly interrupt an ongoing seizure in temporal lobe epilepsy (TLE) mice. Seizures were detected using customized features, such as amplitude, power, regularity, spike width, etc. A binary decision was made based on the combination of the features regarding the onset of a seizure, and accordingly, a command signal was promptly sent to trigger the light. A protocol for a similar configuration as described in ^1^. A flash-FPGA-based system is used in ^53^, where the seizures were detected using an assembly of Morlet wavelet-based decomposition. We integrated the closed-loop optogenetic light delivery and wavelet decomposition module into a single PCB board. The mean detection delay was 575 ms when tested on actual life seizures.

## 3. System configuration and assembly

This section describes the components of our automated closed-loop optogenetics system. The components are (1) online access of data from LabChart, (2) configuration of a deep convolutional regressor, and (3) discharge of TTL pulses for optogenetic light delivery, as shown in Fig. 1.

**Fig. 1.**
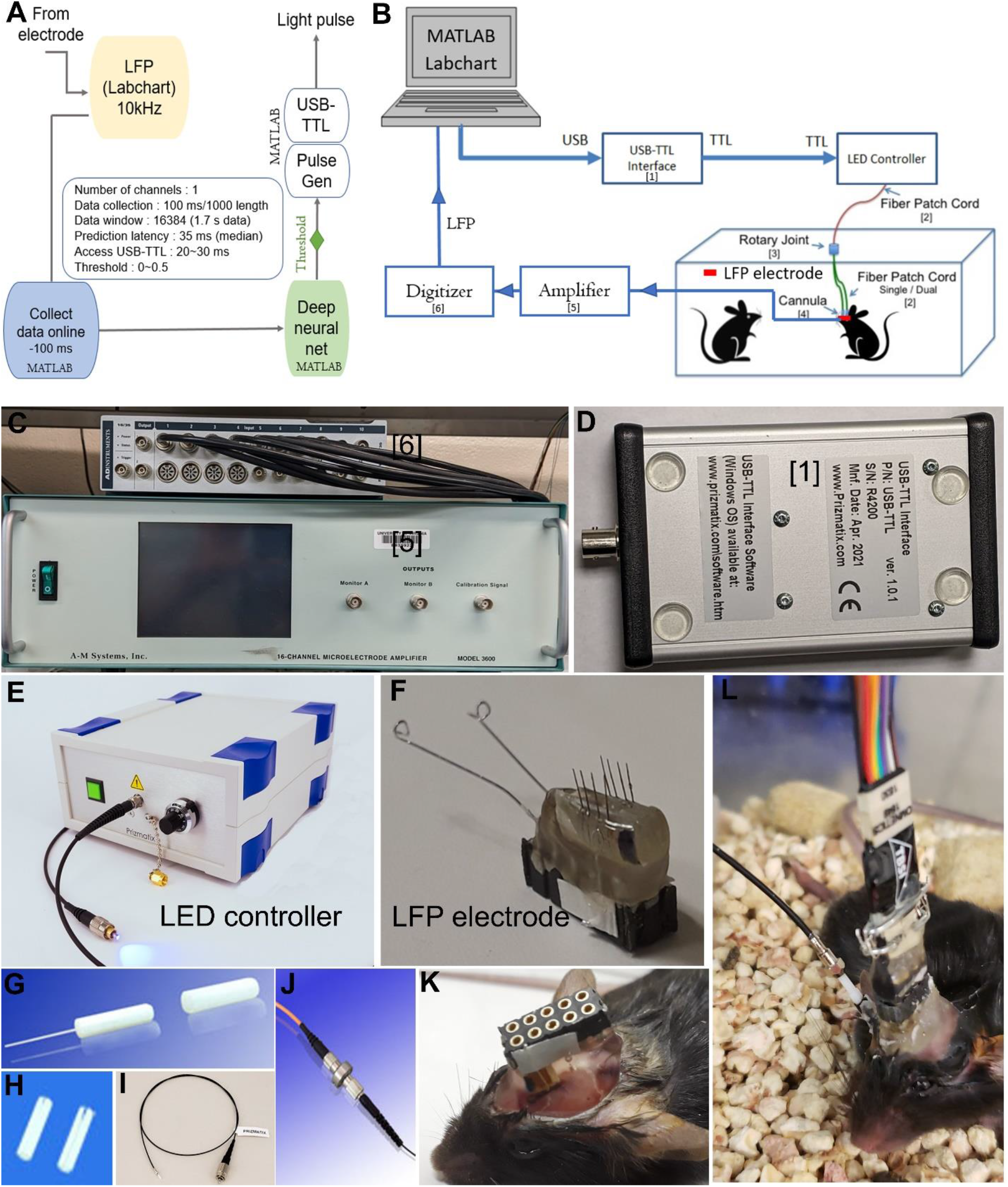
(A) a flow-chart of our single-channel, closed-loop optogenetics setup. The routines for data collection, deep neural net, and sending control signals to the TTL pulse generator are written in MATLAB. (B) The schematics show the connection among the hardware (https://www.prizmatix.com). The assemblies are (C) digitizer and amplifier, (D) USB-TTL interface, (E) LED controller for light delivery, (F) headset with a microelectrode array for LFP recording, (G) cannula with an optical probe, (H) zirconium sleeves for coupling to (I) fiber patch cord, (J) fiber patch cord with rotary joint. (K) a mouse with implanted 10-pin headset containing a customized array of electrodes. It can simultaneously record LFPs from 8 regions. The rest two pins are for the reference and the ground. (L) an example of a mouse with a stereotaxically implanted headset (LFP recording) and an optical fiber (light delivery). The headset and the optical cannula were implanted separately. (E), (G-J) were taken from (https://www.prizmatix.com).

We show a schematic of the closed-loop system in Fig. 1(A). For each mouse, a custom-designed array of electrodes (Fig. 1(F); 60 − 70 *k*Ω resistance of each electrode) with a headset was implanted stereotaxically (Fig. 1(K)-(L)) to record the LFP of neurons over time. The LFP in each channel (or equivalently, each electrode) is amplified 1000 times, passed through a notch filter, and continuously sampled at 10 kHz using a digitizer. The discrete signal is then recorded using LabChart. Along with the LFP recording, the inspection of mouse behavior was performed using video monitoring. The LFP was accessed on the fly using a MATLAB routine, where the routine enabled peripheral communication with the TTL pulse generator Arduino board. We accessed a fixed-length window (let us say L ms) of data in our setup.

The parameters of a pulse are designed using MATLAB function. In our experiments, we used a burst of 5 positive and 5 zero pulses, which are sequentially interleaved, and each pulse has a duration of 50ms, with a total of 500 ms of light delivery, as shown in Fig. 5(A).

The optogenetic toolbox, shown in Fig. 1, contains an LED controller (E), a fiber patch cord (I), a rotary joint, an implantable cannula(G), and ferrules zirconium sleeves(H). The power of LED is controlled via a 10-turn potentiometer (1 – 10 mW/mm^2^) or a 0-5 V analog signal. The rotary joint is a low-friction part designed for freely moving animals. The cannula with desired fiber length was made manually (see the next paragraph). Once the cannula is ready to use, it is tightly coupled to the end of the optical fiber patch cord using a zirconium sleeve.

To prepare a cannula, we first took a fiber with uncleaved ends. We cleaved the fiber to the desired length. The length depends on the depth of penetration of the fiber from the dura. We inserted the cleaved fiber into a ferrule (1.25 mm diameter) and used epoxy to fix the fiber inside the ferrule. The convex end of the ferrule (Fig. 1) was tightly inserted into a puck using a hex key. The puck’s side with the convex side of the ferrule was polished using multiple polishing papers. Finally, we inspected the cannula (assembly of a ferrule and a cleaved fiber) whether a light with a pattern of concentric circles was visible when the cannula was coupled with the fiber patch cord.

## 4. Data description

A seizure is a non-linear rhythmic waveform with time-varying complex morphologies (Fig. 2 (a)). We characterized a cobalt-mouse model of neocortical seizures ^33^. We implanted 1.7 mg of cobalt in the supplementary motor cortex (AP +2.60 mm, ML -1.80 mm; Paxinos Atlas) in the right hemisphere to induce seizures. In this model, the mouse has (1) spontaneous seizures, (2) short-time-interval between consecutive seizures, (3) focal onset electrographic SWD, (4) bursts of beta-gamma oscillations, and (5) suppression of seizures upon administration of phenytoin. The cobalt injury-to the neocortex emulates the cortical injury in human subjects, making this mouse model suitable for studies investigating the neuronal circuitry of cortical seizures.

**Fig. 2.**
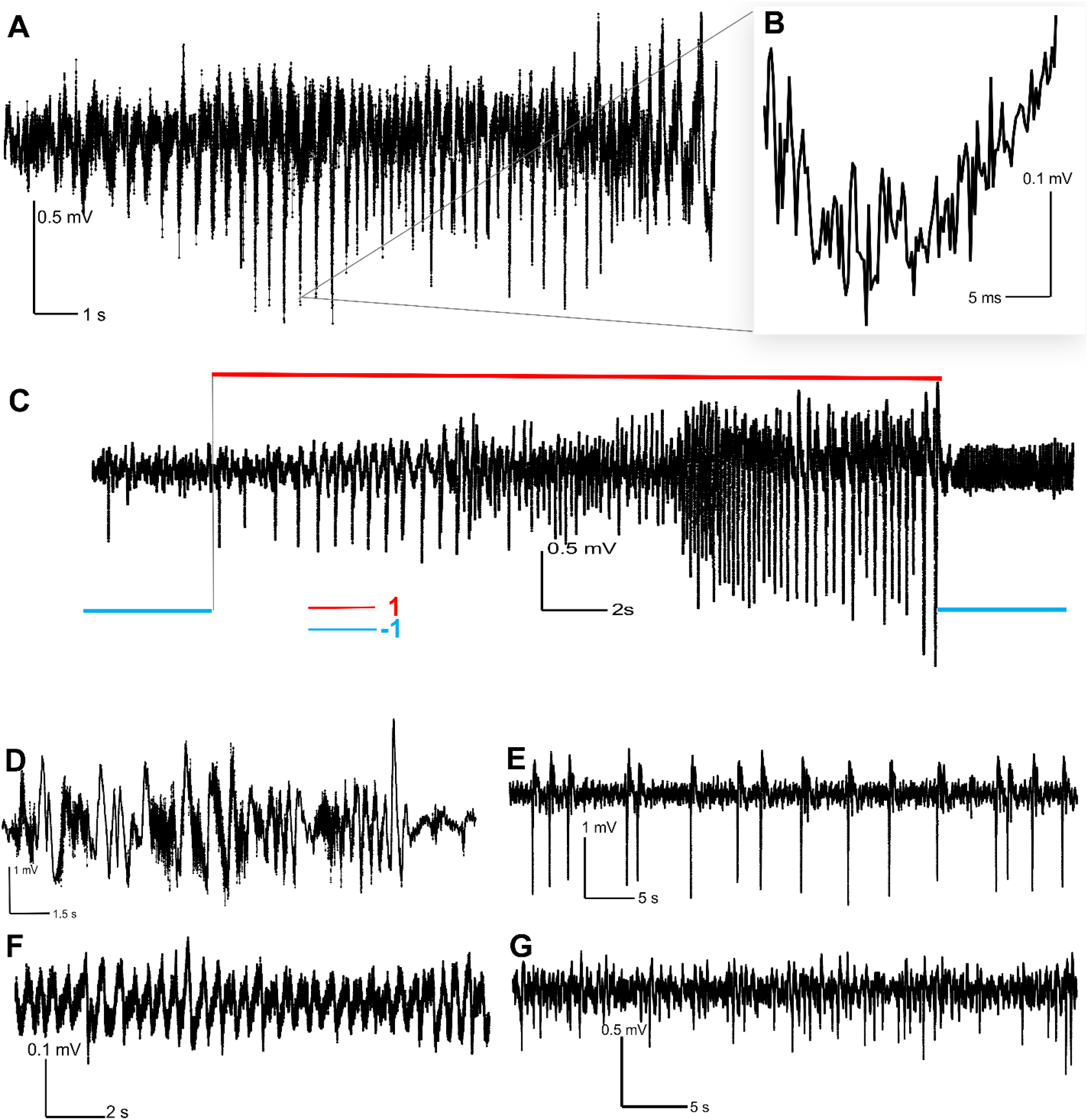
(A) The LFP trace of a seizure, sampled at 10 kHz frequency. (B) A magnified trace near the peak of the seizure (A). The trace shows the high-frequency sampling noise. (C) The LFP trace of a seizure with the corresponding label. The onset of seizure was marked where the magnitude of LFP was at least twice the baseline. For online data, an L (=1000; 100 ms)-length window was constantly accessed with the corresponding label (averaged over L points), preprocessed, and then sent to a deep neural net for evaluation. (D) A noisy baseline with spikes. (E) Epileptiform spikes. (F) a baseline trace of a good signal-to-noise ratio. (G) A baseline with spikes during pre-ictal phase.

We considered 59 seizures (offline) from two groups of mice for training, testing, and validating the performance of our neural net. Mice from the first group (7 mice, five male, and two female) were implanted with cobalt. Mice in the second group (2 mice, 1 male, and one female) were injected with AAV9-CamKII-ArchT-GFP (#99039, Addgene, 120 nl). After viral expression and replication over 14-16 days, we implanted the mice with LFP electrodes and cobalt. We also inserted the optic cannula in the right ventrolateral (VL) nucleus of the thalamus (AP -1.30 mm, ML -2.20 mm, DV -2.80 mm, at 21° such that the probe did not intersect the right lateral ventricle).

Note that CamKII is a promoter-specific to principal glutamatergic neurons. Upon expression of Cre inside neurons, the archaerhodopsin (a polyene chromophore retinal-containing photoreceptor) variant (ArchT; archaerhodopsin from Halorubrum strain TP009) is unlocked and expressed on the plasma membrane. Greenlight (520 nm wavelength) activates ArchT, allowing an outward proton flow, thereby hyperpolarizing the transfected neurons. As a result, the neurons stopped generating and propagating action potentials, leading to neuronal activity suppression. Mice with transfected neurons may differ in seizure morphology, so we did not deliver light to the second group of mice for the first 20 hours during data curation. Out of 59 seizures, we used 39 for training, four for validation, and 16 for offline testing. Once the model was adequately trained and tested with offline data, we deployed it to evaluate its performance on real-time data (5 mice; 26 seizures were interrupted).

We curated an extensive set of control data containing baseline LFP (variable baseline at multiple instances), noise, LFP artifacts due to mouse movements, postictal suppression, burst suppression, epileptiform discharges, and others (see Fig. 2 for details). We extracted 104 files for control data, out of which, 25 files were used for offline training and 13 for validation, and and remaining for testing. The control files contain 8559.2 seconds of data, with 85.59 ± 40.60 s per file.

Before passing the data to the deep neural net, we segmented the data into a set of blocks. It is because, in online mode, we accessed a fixed-length window (let’s say L ms) of data. We experimented using different L values to optimize optogenetic inhibition of neurons (see Results section). In Fig. 1(a), L was set as 100 ms (1000 data points). As briefly mentioned in section 2, the MATLAB routine cannot consistently access 1000 data points. Instead, the accessed data had a variable length with 920 ± 80 data points. The variation rises with the decrease in the length of data access. We maintain a constant length (M = 16384 (= 2^14^)) of data to be passed through the deep learning model as shown in Fig. 1(J). The window length choice depends on desired neuronal activity characteristics. In our case, a typical spike and wave (SWD) discharge during a seizure in rodents has an average frequency of 2 to 3 Hz. With a 10 kHz sampling rate, a window of 16384 data points would contain around 4-5 epileptiform spikes. We assume that the collective morphological features in these 4-5 spikes have sufficient information to classify the window as a seizure or a non-seizure. One may need to set a different window length for other neuronal activities, such as K-complexes and sleep spindles.

The LFP contains high-frequency sampling noise, shown in Fig. 2 (B). To reduce the noise, we preprocessed the data window of length M (=16384 in our case) prior to training and testing the neural net in both offline and online modes. We first used a second-order orthogonal Coiflet wavelet ^21^ with four levels of decomposition, followed by a minimax algorithm to denoise the data window. Next, we applied a running Gaussian window of 100 (∼0.01 *s* of data) length to convolve the wavelet-denoised data window. We followed this procedure for both the control and the seizure data.

We labeled data (+1) if it contained a seizure and (−1) if it did not. The transition from (−1) to (+1) was set at the onset of a seizure, which is defined as *the time instant where the maximum amplitude of the spike (“onset spike”) immediately following the baseline is at least twice the baseline of LFP and the LFP following that time instant corresponds to a spike and wave discharge*. In our work, the transition was set at the beginning of the “onset spike,” not at the peak (Fig. 2 (a)). We defined seizure termination as *the instant where the LFP following a spike returns to the baseline*. The label was set as -1 for all the control data. A deep neural net is data-hungry. To generate a large corpus of data for offline training, we extract 16384 data-window with an offset of 200 samples in case of a seizure. It means that the difference of samples between the start points of two consecutive data *segments* (both of 16384 long) is 200. Therefore, there is an overlap of 16184 data samples between two consecutive data segments. A 200 length skip amounts to 0.02 seconds of data. Roughly, a spike spans 1800 data samples (0.18 s; 10kHz sampling rate). It suggests that a skip of 200 samples does not significantly change the temporal continuity of the data as the input to the neural net. The number of files for control data is substantially larger than the seizure data. The data offset for control data was 800 samples to mitigate the data imbalance effect, which in turn extracts a smaller number of training data from a control file when compared with a file containing a seizure.

## 5. Deep neural net

The set of preprocessed data, each of which is M-length, is passed to six, sequentially-arranged, convolutional filters (Fig. 3). In our model, M is set as 16384 and it is a hyperparameter. Let us assume, *<u>x</u>*(*t*) ∈ ℝ^*M*^ be the input data vector at time t. Let *λ*^*s*^ and *σ*^*s*^ be the pooling (max/average) layer and batch normalization layers, respectively, at *s*^*th*^ stage *s* ∈ {1,2,.., 6}. The corresponding output label is *y*(*t*). The output of our deep model can be given by

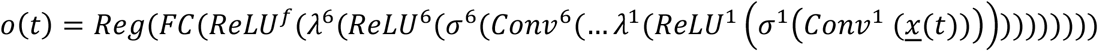

Here, *Reg* denotes the regression layer. If the batch size is b, the regression loss becomes

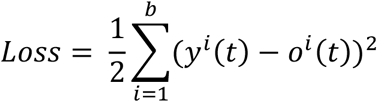

We applied a threshold,*τ*, on the regressed output *o*(*t*) to predict the seizure (label: +1) or no-seizure (label: -1) status of the running window. During training, we used a batch size of 100 samples.

**Fig. 3.**
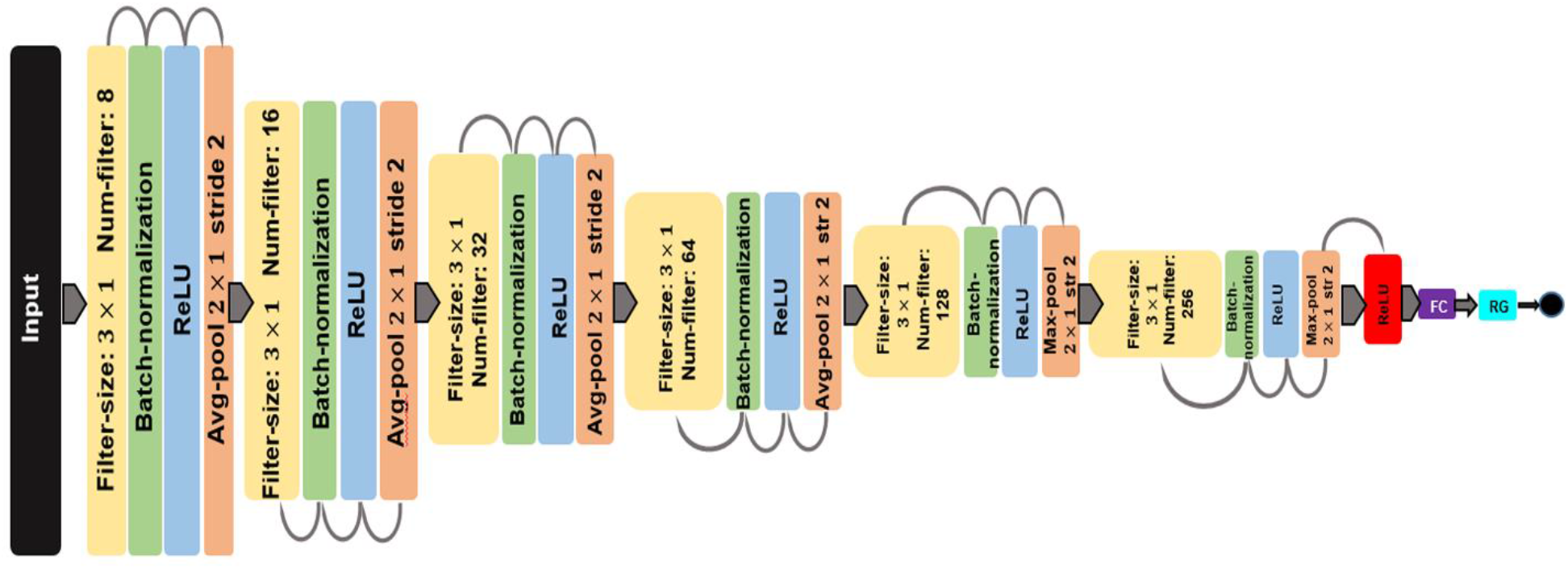
A schematics presentation of a feedforward, sequential convolutional neural network regressor. The input is a vector ∈ ℝ^*M*^. We set M as 16384. ReLU is a rectified linear unit. FC is a fully connected layer. The yellow blocks are convolution filters with specific dimensions as shown. RG is the regression layer.

The neural net training was performed in three stages: batch sizes 6, 8, and 10. The number of epochs in each stage was decided based on (1) whether the performance of the neural net on the validation set is improving or not and (2) the difference between the training loss and validation loss is decreasing (not necessarily monotonically) or not. At first, we randomized the indices of seizures. Let *θ*_*i*_ be the number of segments of the *i*^*th*^ seizure data. Let the total number of segments be 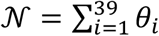 (39 seizures were used for training). The data corresponding to the *i*^*th*^ seizure has a matrix of dimension 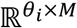. Next, we concatenated the matrix of three seizures, the indices of which were sequentially drawn from the randomized pool. The combined data, 𝒫_*l*_ (*l* ∈ {1,2, …, 13} has a dimension of 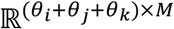, where *i, j and k* are indices from the randomized pool. We randomized the vectors in 𝒫_*l*_ and passed them onto the neural net—this two-step randomization process saved from memory overloading problems during training.

We used Adam as the optimization algorithm for neural network training. The initial learning rate was set as 0.0008. We followed a piecewise learning rate schedule, where the learning rate is dropped by 0.1 after three epochs. At the beginning of each epoch, the data is shuffled once. The tolerance for convergence of neural net’s weights in the training process is set as 10^−5^.

## 6.Results

In this section, we presented the performance efficacy of our closed-loop system in interrupting an ongoing seizure. In literature, the performance of an offline seizure detection algorithm was assessed using a ROC curve. The performance of an online closed-loop system is not only restricted to the detection of seizures. We used a set of measures to evaluate a Type 2 design – latency to of seizure onset detection, duration of pulses after the end of a seizure, early stop of light delivery during a seizure and light delivery during non-seizure LFP.

### Online seizure interruption: optogenetic inhibition

This section shows that our closed-loop system containing the neural net effectively detects and suppresses an ongoing seizure. Fig. 4 (A) shows the green fluorescence protein (GFP) expressed neurons at the right VL thalamus (also called the right motor thalamus), indicating that ArchT receptors were expressed in transfected neurons. The red fluorescence is NeuN immunostaining marking all the neuronal nuclei. The magnified view of the VL thalamus containing the NeuN (red) and GFP (green) fluorescence is shown in Fig. 4(B).

**Fig. 4.**
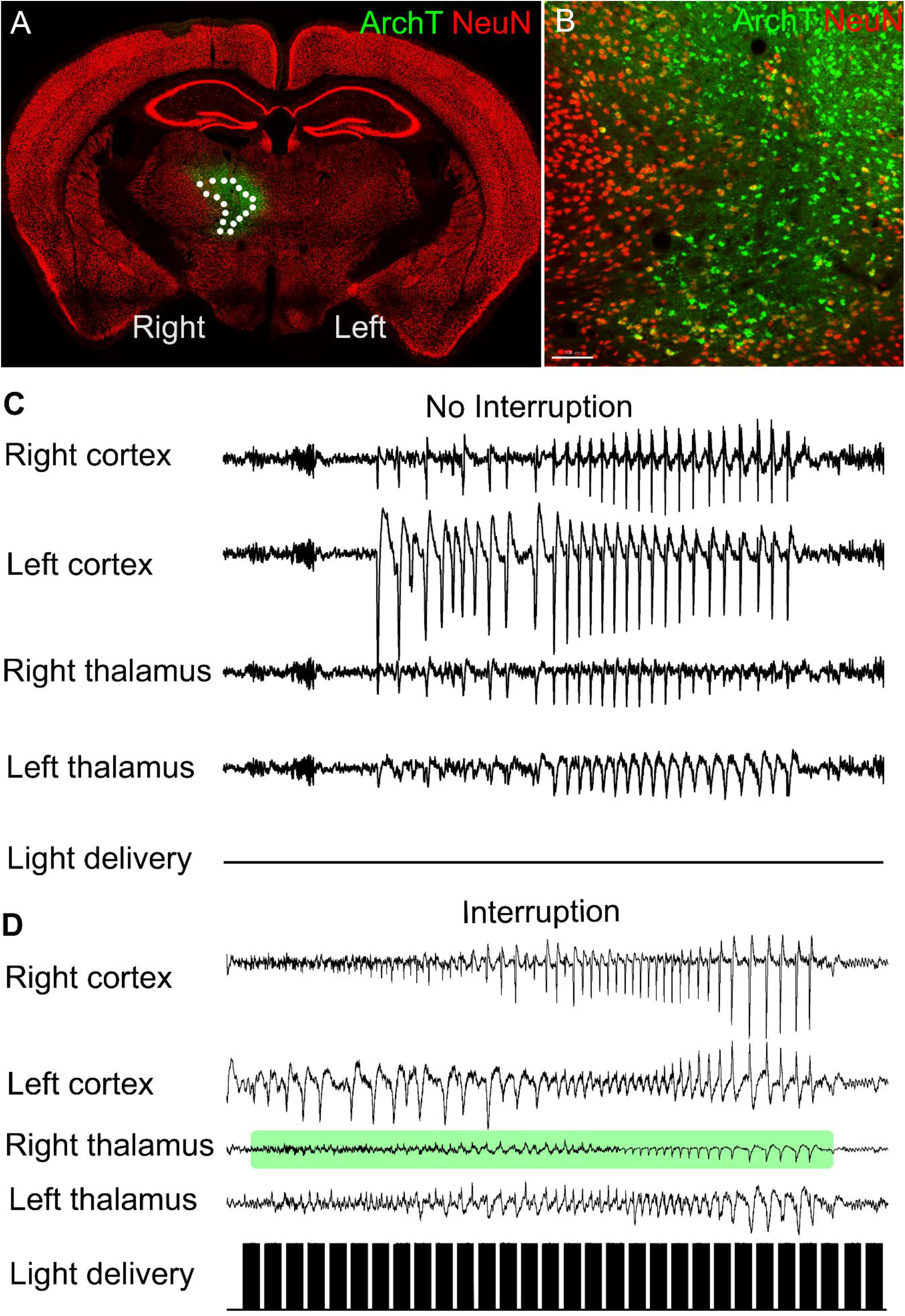
(A) A coronal section of a mouse brain showing the co-expression of NeuN, a marker for neuronal nucleus and GFP (AAV9-FLEX-ArchT-GFP). The green fluorescence protein (GFP) indicates that the cells were transfected and ArchT receptors were expression. The transfection took 14-16 days. The white-dotted region is the right ventrolateral thalamus (VL), which has reciprocal thalamocortical projects with the right motor cortex. (B) magnified VL. The black circle beneath the green ArchT label is the end terminal of the optical probe. (C) The LFP tracing of a seizure at four locations in the absence of light. There was no attenuation/suppression of VL thalamic activity. (D) Suppression of VL thalamic LFP upon light delivery. There was simultaneous suppression at the right cortex.

The LFPs were recorded bilaterally at the primary motor cortex and the VL thalamus. Fig. 4(C) shows the traces of a seizure LFP when we did not deliver light. The seizures at four locations did not exhibit any sign of suppressed neuronal activity. Using the same animal, we delivered light pulses to another ongoing seizure. The neuronal activity was suppressed to baseline upon exposure to the light (Fig. 4(D)). Towards the end of the seizure, there was a rebound neuronal activity at the VL thalamus. This is consistent with the homeostatic principle of neuronal firing, where a group of neurons, which did not express ArchT, showed excitatory activity ^54^. An effective neuronal activity suppression generally precedes this rebound excitation. This rebound excitation can be suppressed by increasing the power of the light. However, we must reduce the neurophysiological damage and possible neuronal death by delivering high-power light.

As mentioned in the Introduction, our deep learning module works on a single-channel recording. We set the LFP at the left motor cortex as the designated channel for online data access because the primary motor cortex has reciprocal excitatory (glutamatergic) projections to the VL thalamus, called the thalamocortical (TC) circuit. Therefore, activity suppression at the right VL thalamus might strongly affect the right motor cortex (Fig. 4(C)). If the right motor cortex was set as the designated channel for data access, a concomitant suppression of neuronal activity caused by the TC suppression would affect the light pulse delivery. We selected the left primary motor cortex because it is largely not suppressed by the right cortical or thalamic suppression.

### Effect of length of online data access (L)(online)

The selection of L is crucial in obtaining a robust suppression of neuronal activity. As mentioned in the Introduction, the strategy employed in our work follows Type 2. In a Type 2 system, a fixed-length (L) data window is accessed for seizure evaluation, and then control signals are sent for light delivery. These two tasks were not performed simultaneously. It suggests that longer L will increase the time difference between consecutive pulses, assuming L corresponds to an ongoing seizure. This delayed, inadequate light delivery contributes to partial activity suppression, as evident in Fig. 5(C). The seizure in this example has a low frequency of arrival of seizure spikes as compared to the ones in (B) and (D). The value of L was set as 1000 (100 ms) in (B) and (D) and 10000 (1s) in (C). For seizures in (B) and (C), we applied the same threshold to the deep neural net output. The one in Fig. 5(B) had a robust suppression. In both of these cases, the light delivery is nearly regular, although the time delay between the arrivals of consecutive pulses differs. For (B), the time difference between consecutive pulses (assuming that two consecutive windows were evaluated with +1: seizure) is 157 ms. For (C), the time difference between consecutive pulses (assuming that two consecutive windows were tagged with +1: seizure) is 1053 ms. Another point worth noticing is the onset detection delay of a seizure. In case of L = 1000 (Fig. 5(C)), the detection delay is considerably larger than with L = 100 in Fig. 5(B).

**Fig. 5.**
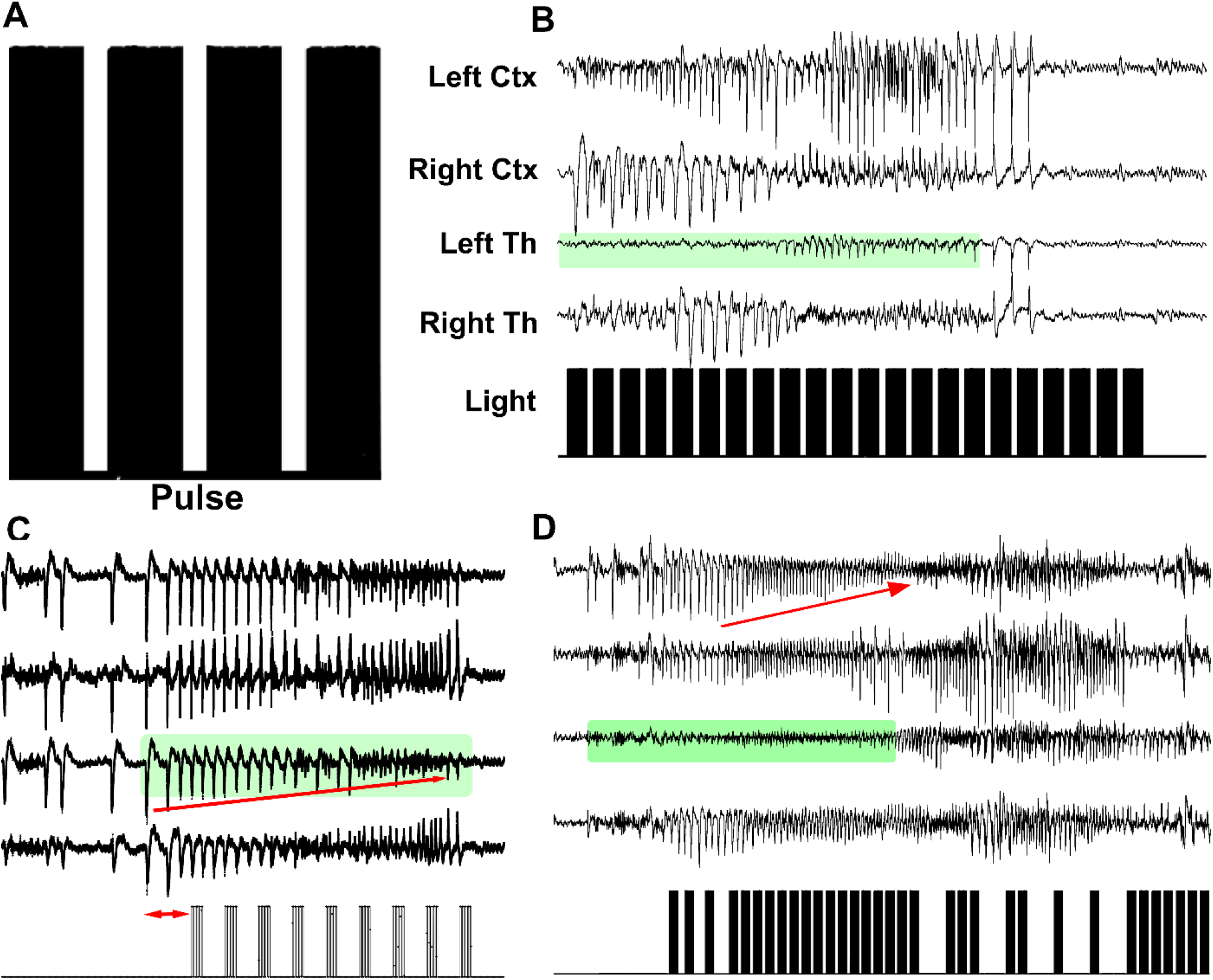
(A) A pulse train. Each thick black column contains five pulses of 50 ms duration each. Each column is 500 ms in duration. (B) The LFP trace of a seizure where light delivery suppressed the VL thalamic activity. The latency of seizure onset detection was 130 ms. The light delivery continued after the seizure ended (postictal suppression phase). L was set as 1000 (100 ms). (C) The regular but wide-spaced light probing. Here L was set as 10000 (1s). The light was still able to induce suppression but not as effective as in (A). The latency of the neural net to detect the seizure onset was also larger compared to (A). (D) The LFP trace of seizure. L was set at 1000. The threshold was *τ* = 0.8. A high threshold rendered the light pattern irregular, resulting in reduced suppression. In addition, the latency of the neural net to detect the seizure onset was also large. The red arrow indicates cortical suppression.

### Effect of threshold (τ) of deep neural net output (online)

As noted in section 4, we applied a hard threshold to the output of our deep neural net regressor. An output below *τ* was no-seizure (−1). An output above *τ* was a seizure (− 1). An ideal choice of *τ* would be 0. However, we empirically found a range (0.1 – 0.35) performing well for our purpose. We set the threshold for the seizures in Fig. 5(B) and (C) as 0.15 and for (D) as 1.0. Generally, lowering *τ* will increase false positives (i.e. light delivery during no-seizure LFP). Such action will also prompt continuing light delivery after a seizure ends, negatively affecting cellular physiology. However, it decreases the latency to detect the onset of seizures. In addition, the light delivery is regular during an ongoing seizure. On the other hand, increasing *τ* will have the opposite effects. Noteworthy among them is the irregular pattern of light delivery during a seizure, as can be observed in Fig. 5(D).

In contrast to the examples in Fig. 5 (B) and (C), where we set the left (contralateral) motor cortex as our designated location for single-channel data access, we set the right (ipsilateral) motor cortex as the designated one in Fig. 5(D). In Fig. 5(D), neural activity was suppressed during a regular delivery of light (green shadowed region). This was followed by a brief rebound excitation, during which three pulse trains of light were launched. The three pulses could not reduce this rebound activity at the right thalamus, possibly because we employed low power of light (∼4mW) to reduce cellular damage. Also, note that consistent with the TC circuit, the right motor cortex had suppression of neuronal activity. In contrast, the contralateral left motor cortex LFP did not have a sign of suppression. The light delivery briefly stopped when the LFP at the right motor cortex reached the baseline. At this point, the deep neural net output scores were lying in (0, 0.4) and thus were unable to trigger the light because of the high threshold *τ* = 0.8. Following the cortical suppression and stoppage of light delivery, the seizure amplitude at the left motor cortex was elevated. At this instant, the thalamic neurons were not entirely suppressed. From this stage onward, higher *τ* (>0.4) precludes a regular light delivery. This irregularity could not effectively suppress the thalamic activity anymore. It indicates that the activity suppression of a neuronal population is a gradual process which has a finite time constant for inactivation. Such a process requires nearly continuous light delivery (For details, please see the Discussion).

### Quality of seizure interruption

[Offline] our deep neural net effectively determines the control for light delivery. It produces outputs (regressed scores) that do not send signals to deliver light for control outputs. The output is resilient to various control waveforms, as shown in Fig. 6. Fig 6 (A) shows the neural net output as “PredictedLabel”. The output scores were below the threshold (dotted line) during the pre-ictal and mostly postictal phases. The output scores reside in (−1, +1). Interestingly, as the seizure frequency waned midway, the neural net output scores dropped, possibly because the data window started containing fewer spikes. This is also true towards the end of the seizure in Fig. 6(A).

**Fig. 6.**
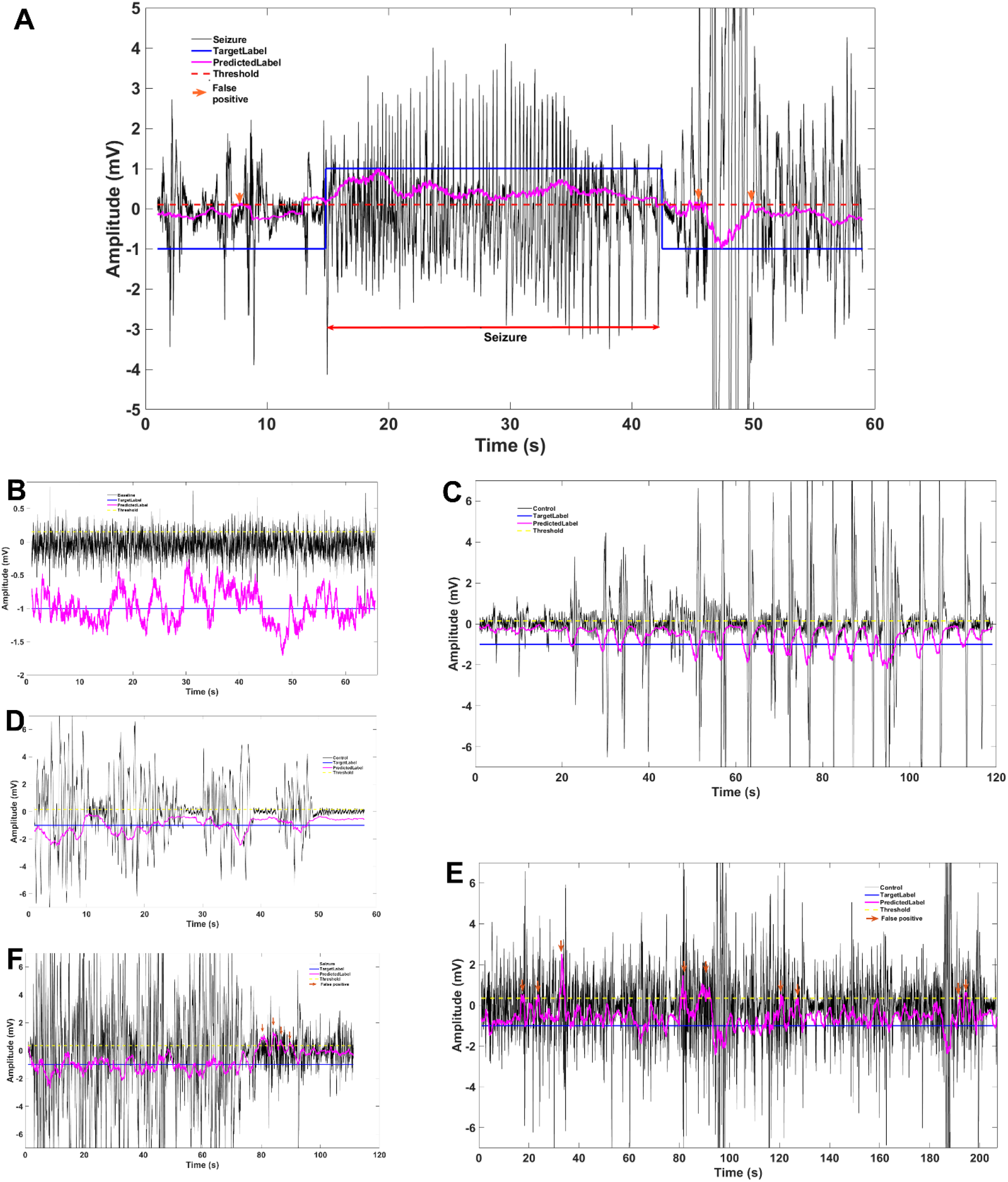
Performance of our neural net using offline data. TargetLabel is the desired output of our neural net (M = 16384, L = 1000). PredictedLabel is the output by our neural net. False-positive detections are shown in downfaced arrows. The threshold, *τ* was marked with a dashed line in each plot. (A) Performance of our neural net on a seizure LFP. The latency to detect the seizure onset was 80 ms. There were three false positives during the preictal and postictal phases. (B) - (E) Performance of our neural net is also shown in several control data.

The neural net outputs for a baseline are shown in Fig. 6(B). The baseline magnitude roughly varied from (−0.5, 0.5). The predicted label showed an oscillatory behavior around the target label, -1. Fig. 6(C) showed the neural net outputs for the LFP of epileptiform discharges / large spikes. There was no false-positive detection by our neural net. Note that neural net scores dropped at the onset of each large spike and exhibited a rebound at the end of the spike. This showed that the neural net responses to data window (16384-length) containing a single spike were strongly guided towards - 1. Fig. 6(D)-(E) showed the neural net responses in cases of occasional noises and artifacts, possibly stemming from mice movements. The neural net did not yield false-positive detections. Fig. 6(F) presented an LFP recording containing significant noise. The neural net performed well on the noisy signal, yielding output close to -1. However, the transition from the noise to a spikes level elevated the neural net responses. Such elevation prompted a short burst of light delivery (false positives). Fig. 6(G) exhibited LFP recordings of prolonged duration of such noisy baseline activity and the corresponding neural net response over time.

[Online] we report the performance of our closed-loop optogenetics setup in vivo in Table 1. The LFP of each mouse was recorded for 40 hours. We activated the closed-loop module for 20 hours for each mouse. The light was turned off for the rest of the 20 hours, and it acted as the control for the same mouse. We ruled out any subject-wise differences between controls and experiments in the performance of our model. The desirable range of *τ* was [0.15, 0.35] for in-vivo experiments. Please note that *τ* was set as zero for offline training of our neural net. The elevation of *τ* for in-vivo from 0 to [0.15, 0.35] indicates that there were variations in LFP during in-vivo experiments that were unaccounted during offline training. A higher threshold (animal 3) for noisy baseline lowered the false positives (FP ratio). However, the latency to seizure onset detection was increased. In a set of seizures, a higher threshold stopped the light delivery before a seizure also ended. In addition, higher threshold rendered the light pulse delivery irregular in time (Fig. 5(D)). For noisy baseline (animal 1), the neural net successfully detected seizure onset (120 ms in mean), and regular light delivery. However, the FP ratio was elevated.

**Table 1:** The analysis of the performance of our closed-loop optogenetics set up when it was deployed for in-vivo experiments. Pulses postictal is the duration of how long the light was delivered immediately after a seizure ended. FP-ratio is the ratio of the duration of pulses (excluding the pulses during seizures) to the total observed duration. Latency-onset-light is the latency of seizure onset detection. In total, 26 seizures were interrupted.

| Filename | $\tau$ | SZ duration<br>(s) | Pulses<br>post-<br>ictal (s) | Latency-onset-light<br>(ms) | FP-ratio | Light<br>Trigger<br>(hrs) | Comment |
| --- | --- | --- | --- | --- | --- | --- | --- |
| Animal -1 | 0.15 | 31.06 ± 11.71 | 3 ± 1.03 | 120 ± 37.3 | 0.077 | 20 | Noisy baseline |
| Animal - 2 | 0.35 | 18.83 ± 6.33 | 4.08 ± 4.04 | 1833.3 ± 2309.4, Median = 500 | 0.011 | 20 | Baseline in (-0.5, 0.5) mV. Significant movement artifacts |
| Animal - 3 | 0.8 | 24.2 ± 4.8 | 0.2 ± 0.27 | 3500 ± 1008.5 | 0.0017 | 20 | Premature termination of light delivery in 2 seizures |
| Animal - 4 | 0.35 | 49.83 ± 11.47 | 0.92 ± 0.58 | 200 ± 489.8 | 0.0027 | 20 | Baseline in (-0.5, 0.5) mV. |
| Animal - 5 | 0.15 | 22.5 ± 3.69 | 1 ± 0.707 | 255 ± 206.15 | 0.0006 | 20 | Baseline in (-0.5, 0.5) mV. |

### Effect of fixed window length, M [offline]

The quality of suppression depends on the choice of M. Fig. 7(A) shows the neural net output scores where M, the window length, was set as 8192. The sampling frequency was 10 kHz. In general, an 8192-length window contains, on average, two spikes in an 8192 length LFP during an ongoing seizure. In contrast, a 16384-length window contains on average 5 spikes during a continuous seizure. When M is 8192, the neural net generates a number of false positives, as can be observed in Fig. 7(A) (the predicted label is below the threshold). It is interesting to note the neural net scores during the onset and end of the seizure. It may be because the frequency of spikes increased during the middle of the seizure and the neural net, trained on M = 8192, failed to recognize it. A 16384-length window encompasses a large variation in spikes compared to an 8192-length window. In short, a 16384-length data is comprehensive coverage of information of spikes than an 8192-length data. So, neural nets trained on a 16384-length are trained robustly.

**Fig. 7.**
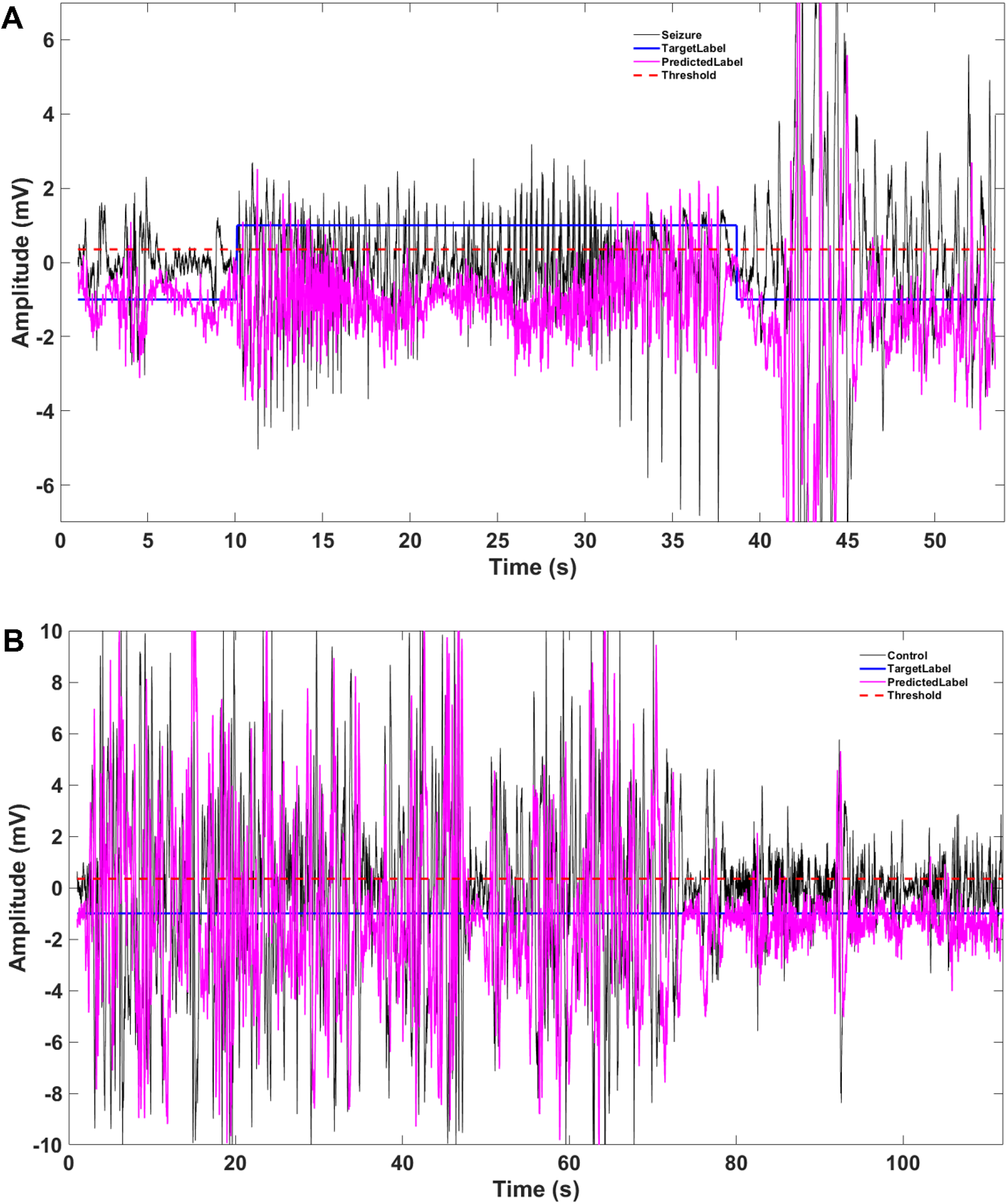
Performance of neural net trained with M = 8192 windows of data on (A) a seizure and (B) a noisy control data. The performance of our neural net (M = 16384) on the same control data is shown in Fig. 6 (F).

Another important point is that neural net scores (M = 8192) appeared to adopt the morphology of individual spikes at the onset and end of the seizures (Fig. 7(A)-(B)). Such behavior was not observed in the case of a neural net with M as 16384. It is primarily because the proportion of newly added data to M for two cases was 0.06 (= 1000/16384) and 0.122 (= 8192/16384). Therefore, the 16384-length window has much less newly-added temporal context, thus inducing smoothness in its output scores. Fig. 7(B) shows the neural net (M = 8192) responses for the LFP for which we also provided neural net (M = 16384) scores in Fig. 6(F). It is evident that neural net (M = 8192) scores are sensitive to noise and generate many false positives.

### Comparison with state of the art

We evaluated the performance of the state of the arts in offline mode and demonstrated the superior performance of our method (see Table 2). The performance metrics include seizure detection accuracy, median latency of seizure onset detection, post-ictal continuation of light delivery. We added comments to each method that we implemented to compare against our method.

**Table 2:**
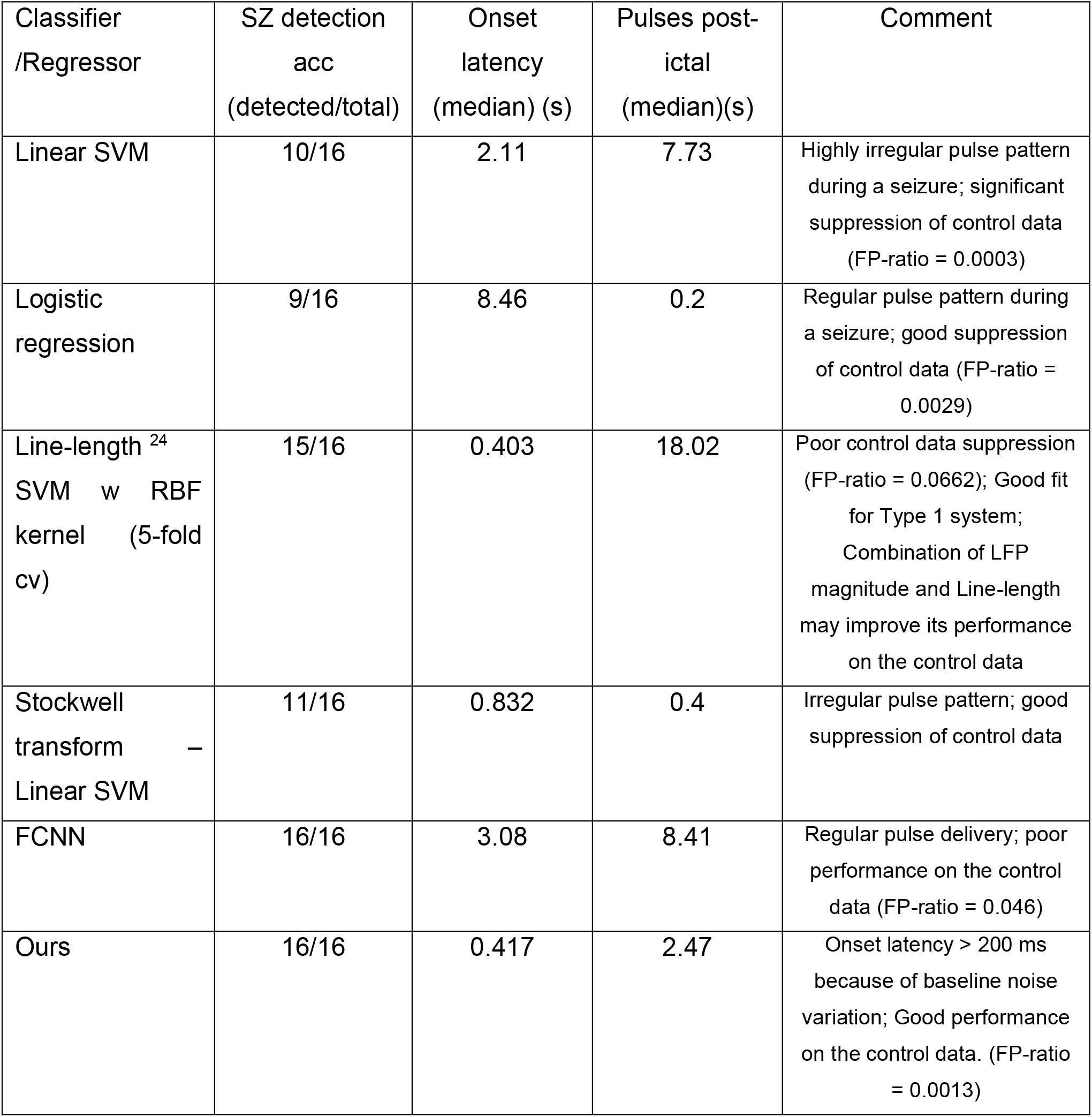
The comparison of performance using different measures between the state of the arts and our method. FP-ratio is the ratio of the duration of pulses (excluding the pulses during seizures) to the total observed duration.

| Classifier /Regressor | SZ detection acc<br>(detected/total) | Onset latency<br>(median) (s) | Pulses post-ictal<br>(median)(s) | Comment |
| --- | --- | --- | --- | --- |
| Linear SVM | 10/16 | 2.11 | 7.73 | Highly irregular pulse pattern during a seizure; significant suppression of control data (FP-ratio = 0.0003) |
| Logistic regression | 9/16 | 8.46 | 0.2 | Regular pulse pattern during a seizure; good suppression of control data (FP-ratio = 0.0029) |
| Line-length<sup>24</sup> SVM w RBF kernel (5-fold cv) | 15/16 | 0.403 | 18.02 | Poor control data suppression (FP-ratio = 0.0662); Good fit for Type 1 system; Combination of LFP magnitude and Line-length may improve its performance on the control data |
| Stockwell transform – Linear SVM | 11/16 | 0.832 | 0.4 | Irregular pulse pattern; good suppression of control data |
| FCNN | 16/16 | 3.08 | 8.41 | Regular pulse delivery; poor performance on the control data (FP-ratio = 0.046) |
| Ours | 16/16 | 0.417 | 2.47 | Onset latency > 200 ms because of baseline noise variation; Good performance on the control data. (FP-ratio = 0.0013) |

For evaluation, we arranged the LFP data in blocks, where each block has a length of 16384. The corresponding output labels were obtained as real values in [0, 1]. If the model that we compared with our method is a classifier, we binarized the output label as 0 or 1. Using raw LFP data ∈ *R*^*No*−*of*−*block*×16384^, we trained an SVM model, a logistic regression model and an FCNN. We extracted features from each block if needed. For example, we extracted line length^24^ and Stockwell transform^55^, and evaluated the performance of each of them. In case of the Stockwell transform, a single time series outputs a 2D map consisting of the frequency and time information. We computed the power of the S-transform in the frequencies in [4 Hz,40 Hz], discarding the power of the signal in the delta frequency range ([0.1 Hz, 4 Hz]). In addition, we implemented a fully connected neural network regressor (FCNNR) with 4 hidden layers and one input and one output layer (Input (16384) -> hidden1 (2048) -> hidden2 (256) -> hidden3 (32) -> hidden4 (4) -> output (1)). It is reported in ^45^ that CNN performed better than an FCNN. We found the same evidence in our offline experiments.

For 16384 dimensional data, we used linear SVM. Each classifier was 5-fold cross-validated and the regularization strength Lambda was searched in the interval of [10^−6^, 10^0.5^]. We checked each cross-validated model and their predicted scores for each Lambda value. We reported the result of the model that achieved the highest true positive and true negative rates.

## 7. Discussion

We propose a flexible and modular design of a closed-loop optogenetics system with a deep neural net for seizure detection. Conventional state-of-the-art systems for closed-loop optogenetics have limitations related to flexibility, scalability, and feature-detection algorithms^16^.

Many existing systems use a customized headset for simultaneous LFP recording and optical stimulation. The LFP/EEG headsets and optical probes are implanted separately in our case. Another problem stems from implementing algorithms to recognize “desired” neuronal activity. For instance, in our experiments, the seizure was the desired neuronal activity. We expected the algorithm to detect each seizure while ignoring other neuronal activities and noise in LFP. The detection and prediction of a neuronal activity encompass a broad class of problems – activity onset detection, activity detection, and early activity prediction, detection of activity termination. The selection and configuration of a machine learning model depend on the problem. A closed-loop system should have the flexibility to incorporate a series of such functional models and provide a way for a user to select the models that the user wants. Hardware-based implementation of a detection algorithm for a specific activity does not have the flexibility to include additional functional modules. We have software-based implantation of the neuronal activity detection algorithm, where other modules can be incorporated easily. The settings can be also be extended to incorporate multichannel LFP, where the LFP of each channel is evaluated using the same deep network. Trained models of desired activities other than seizures can be seamlessly integrated into our system.

The selection of opsin is a crucial factor. In some studies, halorhodopsin (NpHR: a chloride pump derived from the halobacterium) was used, which sends chloride ions into the cell causing hyperpolarization ^2^. Trafficking chloride ions into neurons may alter physiology and induce unwanted effects (for example, reducing GABAergic inhibition) when the cells are exposed to sustained light delivery ^56^. ArchT is a proton pump, ejecting H^+^ from cytosol to the extracellular space. It may cause pH-dependent activation/block of N-methyl-D-aspartate receptor (NMDAR) subunits^57,58^. Higher than the normal extracellular pH can elevate NMDAR activity, promoting the firing of un-transfected neurons in the neighborhood of the transfected cells. So prolonged exposure of neurons expressing ArchT to green light can also affect cellular physiology. The protocol for light delivery must follow the kinetics of the selected opsin.

The median time for ArchT expressing neurons from the light trigger to the onset of the decrease in firing rate was 60 ms ^59^. After the cessation of light delivery, ArchT expressing neurons recovered to the baseline firing rate within 740 ms (median) and they did not show rebound elevation of firing rate after returning to the baseline ^59^. A pulse (+1: light ON) of 50 ms for light delivery would sufficiently hyperpolarize the transfected neuron. The next pulse (0: light OFF) would curb the outflow of H+ to the extracellular matrix as ArchT receptor is an outward proton pump. At this stage, the neuron is still hyperpolarized (assuming the recovery curve is linear right after the light is off, the neuron is still 50/740 ≈ 7% recovered). This hyperpolarized neuron then experienced the next 50 ms-pulse (+1: light ON) of light delivery. After five pulses of (+1: light ON) and five pulses of (0: light OFF), our closed-loop settings fetched the next L = 1000 (100 ms) length online data during which light delivery was in the OFF state. The median time of data access, preprocessing, neural net output generation, USB-TTL access, and the trigger, put all together, is 175 ms. Overall, the transfected and hyperpolarized neurons had 225 ms (175+50 (the last 0-pulse): ≈ 30% recovered) for recovery before the next pulse train of light hit the neuron. This design protocol renders the pulse train nearly continuous, curbs the accumulation of H^+^ in the extracellular domain, precludes the necessity of high-power light delivery, and maintains the hyperpolarized state of transfected neurons during a seizure.

We sampled LFP at 10kHz. A change in the sampling frequency will affect the performance of our neural net. Let us assume a recording system with a sampling frequency of 500 Hz. Online access of L = 50 (100 ms) length data would be appended to the running window, M. M can be set as 1024 (it is the number that can be expressed as an integer power of 2, which is immediately larger than 850 or 1.7s of data at 500 Hz). With such settings, the neural net can be trained from the beginning. Alternatively, L = 50 can be added to the existing settings of M (running window for seizure evaluation by the neural net) with M as 16384. With training data sampled at 500 Hz (the user is required to curate), the FC (fully-connected) layer in Fig. 3 can be re-trained using our pretrained model.

## Animal guidelines

We used five C57Bl6 mice (3 male and 2 female; 8-12 weeks old) for in vivo experiments. For training the neural net, we used nine C57Bl/6 mice (6 male and 3 female; 8-12 weeks old). All experiments and procedures were performed under the guidelines set by the University of Virginia Animal Care and Use Committee.

## Data availability

Data are made publicly available in github (https://github.com/50-Cent/Closed-loop-optogenetics-KapurLab).

## Code availability

This code is made publicly available in github (https://github.com/50-Cent/Closed-loop-optogenetics-KapurLab).

## Acknowledgements

Grants from the National Institute of Health (RO1 NS040337, RO1 NS044370 to J.K.) and the UVA Brain Institute supported this work.

